# Reconstitution in JCVI-syn3B suggests the roles of the *Spiroplasma* Fibril protein (Sfibril) in swimming

**DOI:** 10.64898/2026.11.17.765575

**Authors:** Ali Ahsan, Hana Kiyama, Yuhei O Tahara, Yuya Sasajima, Makoto Miyata

## Abstract

*Spiroplasma*, wall-less helical bacteria of the class Mollicutes, swim without flagella by shifting the handedness of their helical body. This motility is driven by an internal ribbon composed of actin-superfamily MreB proteins (SMreB) and the *Spiroplasma*-specific Fibril protein (Sfibril), which accounts for nearly half of the ribbon mass. However, because helical shape and swimming have been reconstituted with SMreBs alone, the role of Sfibril has remained unclear. Here, we expressed Fibril from *Spiroplasma eriocheiris* under its native promoter in the minimal synthetic bacterium JCVI-syn3B (syn3), in which swimming had been reconstituted with SMreB proteins, and analyzed cell morphology and swimming. Sfibril formed filaments with chained ring units in syn3 cells and, by itself, induced cell elongation and left-handed helical shape. Elongated cells expressing Sfibril recovered their length after hypo-osmotic shock, indicating that Sfibril confers morphological robustness. In cells expressing SMreB4 and SMreB5 from *Spiroplasma*, Sfibril increased the fraction of helical and motile cells, extended the distance of continuous kink propagation, reduced cell bending, and increased traveling distance. In addition, Sfibril altered the handedness of SMreB5-driven helices from exclusively right-handed to a mixture of right- and left-handed. These results suggest that Sfibril is not essential for helicity shift but stabilizes the cell body and supports efficient, processive helicity shifts.

**IMPORTANCE:** *Spiroplasma* are helical bacteria that swim by shifting their helical handedness. This shift is caused by a “ribbon structure” in the bacterial cell. The shift of the ribbon structure is driven by MreB proteins, which are related to actin, the major component of thin filaments in muscle. In this study, we focused on “Fibril protein”—another major component of the ribbon structure—and showed that this protein supports the smooth helicity shift like a “flexible shaft”, through reconstitution in a minimal synthetic bacterium.

**Graphical summary:** 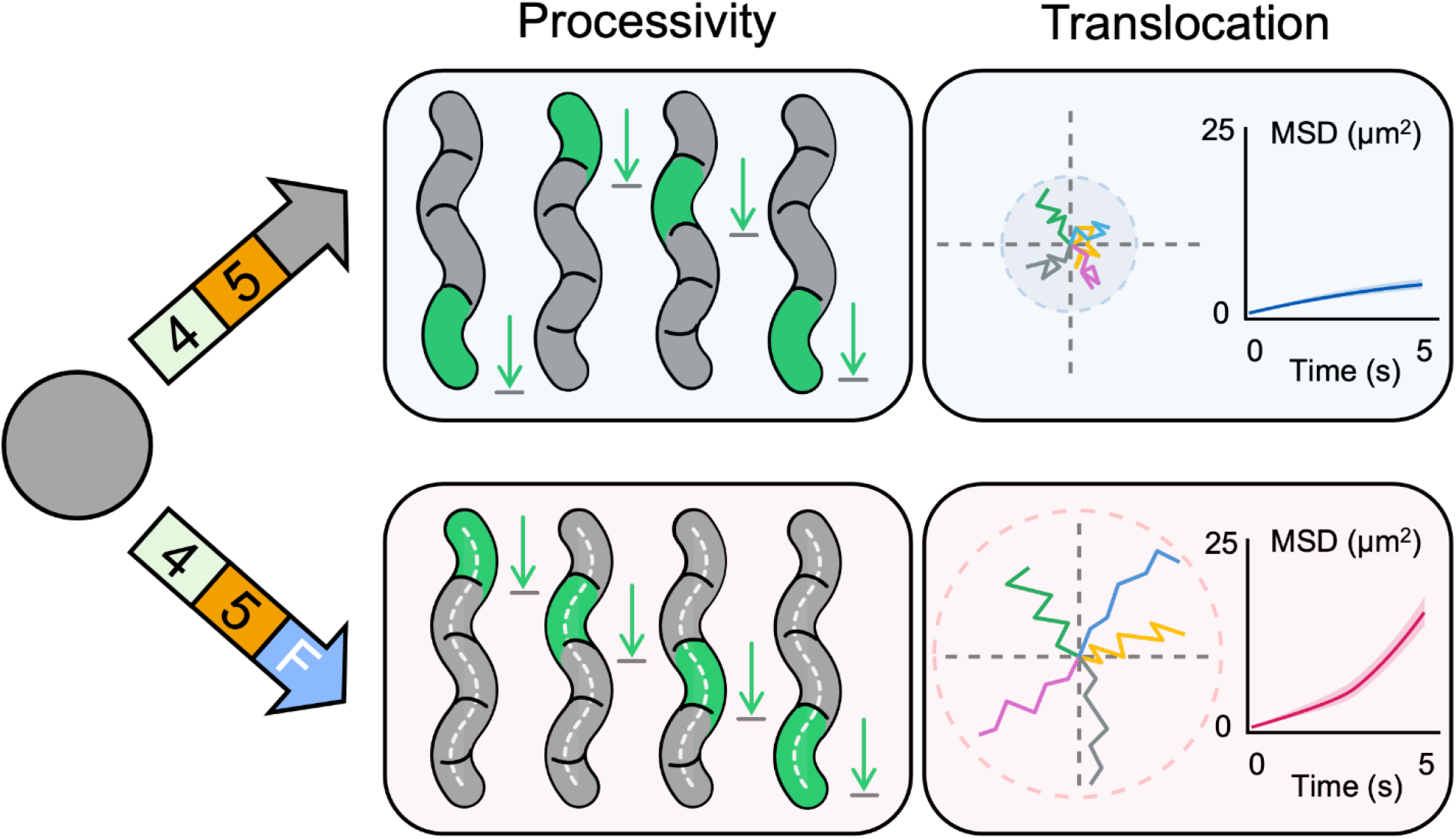

## INTRODUCTION

Helical bacteria of the genus *Spiroplasma* (class Mollicutes) lack flagella and pili, yet execute rapid swimming through viscous media (1–4). This motility is driven by helical shifts between right-handed (RH) and left-handed (LH) geometries. The boundary between these opposing helical segments, termed “kink,” propagates processively along the cell body. The cell body behind the kink rotates, pushing water backward (5–9). Ultrastructural studies suggested that the helicity shift is driven by an internal cytoskeletal structure termed the “ribbon” that runs along the shortest innermost line of cell curvature. This ribbon consists of actin-superfamily MreB proteins (SMreBs) and the *Spiroplasma*-specific ∼60-kDa Fibril protein (Sfibril) (1, 8, 10–14). Sfibril accounts for roughly 10% of total cellular protein and nearly half of the ribbon mass (15–17). The single-stranded Sfibril filament isolated from *Spiroplasma eriocheiris* consists of rings (11 nm wide, 6 nm long) connected by a backbone cylinder with an 8.7-nm interval and a twist along the filament axis (18, 19). The amino acid sequence of 512 residues is divided into two domains. The N-terminal domain (residues 1–228) shows a structure similar to that of S-adenosylhomocysteine nucleosidase, while the C-terminal domain (residues 229–512) does not show similarity to other proteins (5, 18–20).

For decades, Sfibril was thought to be a contractile linear motor driving helicity shift through length changes coordinated with adjacent protofilaments (11, 12, 21, 22). However, this paradigm was challenged by two findings: (i) A nonsense mutation in the gene for class 5 MreB of *Spiroplasma* (SMreB5 for short) abolished both helical shape and swimming motility (5, 6, 23); (ii) *Spiroplasma* motility was reconstituted in JCVI-syn3B (syn3 for short) by expressing a pair of SMreB4 and SMreB5 (24). What, then, is the role of Sfibril in *Spiroplasma* cells?

syn3 is a minimal bacterium possessing a genome that encodes only 473 genes essential for life (25, 26). The genome was designed based on *Mycoplasma mycoides*, a relative of *Spiroplasma*, chemically synthesized, and then transplanted into *Mycoplasma capricolum* cells. syn3 is useful for examining gene functions, especially for genes from related species. In a previous study, our group found that a pair of SMreB4 and SMreB5 from *Spiroplasma eriocheiris* can reconstitute *Spiroplasma*-like motility, and both SMreB5 and Sfibril can cause cell elongation and helical cell shape (24, 27–29).

In this study, we characterized syn3 cells expressing Sfibril with SMreB4 and SMreB5, and suggest that Sfibril contributes to the robustness and efficiency of swimming.

## RESULTS

### Reconstitution and structural characterization of Sfibril in *JCVI-syn3B*

To investigate the cellular role of Sfibril, we expressed the *fibril* gene from *S. eriocheiris* in syn3 (Fig. 1 and Fig. S1). The DNA fragment including the native promoter was inserted into the *loxP* locus of syn3 genome (Fig. 1A). SDS-PAGE and peptide mass fingerprinting (PMF) showed expression of Sfibril at the position expected from its calculated mass (58.7 kDa) (Fig. 1B; Table S1). The amount of Sfibril was 2.5-fold higher than that obtained with the pTuf promoter in our previous study (24).

**FIG 1.**
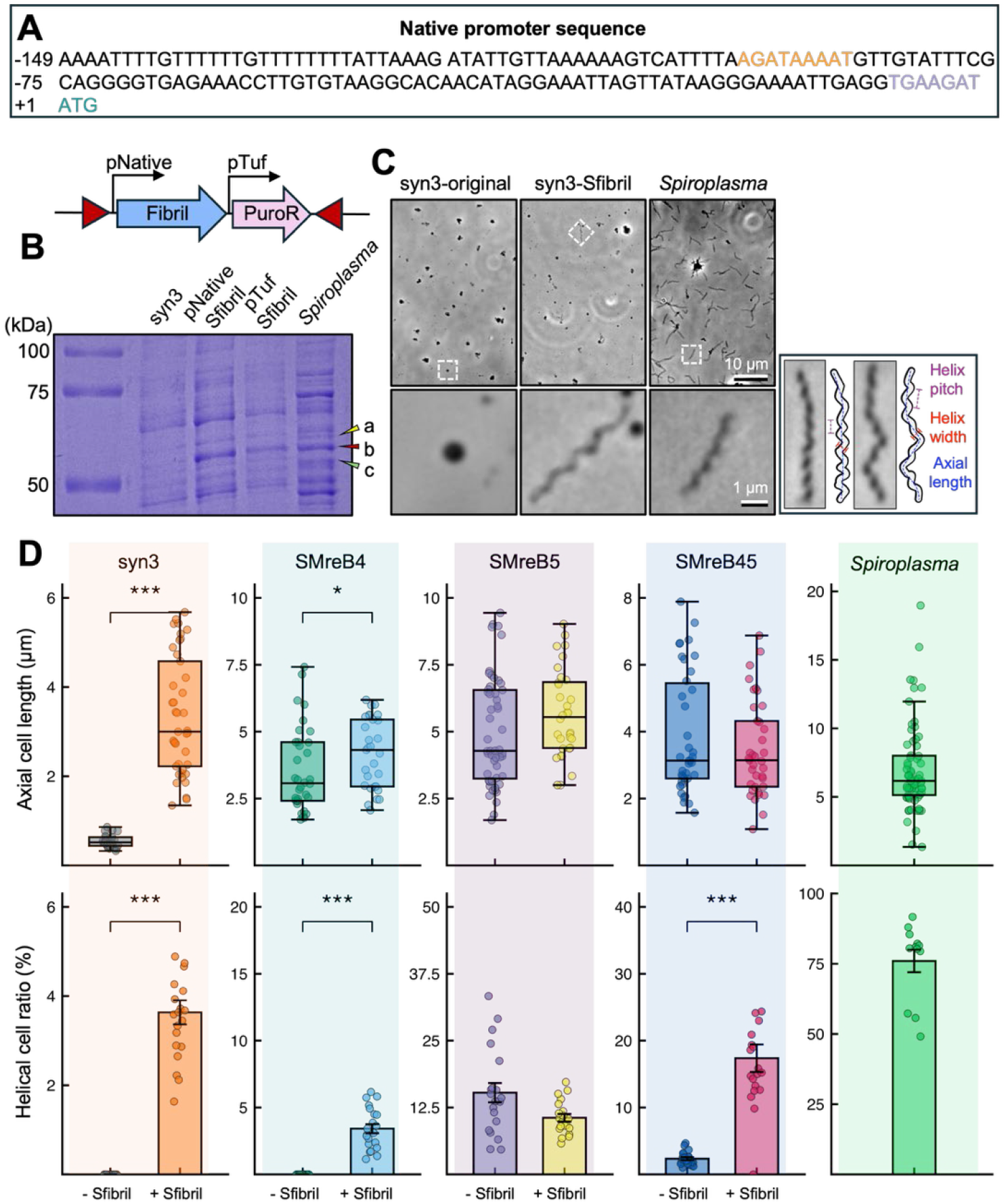
Effect of Sfibril on elongation of syn3 cells. (A) DNA fragment inserted into *loxP* locus of syn3 genome. The fragment contains the *S. eriocheiris fibril* gene and preceding sequence including the native promoter (orange and purple colored). The initiation codon is colored green. Red triangles indicate the *loxP* recombination sites. (B) SDS-PAGE analysis of cell lysates. Protein bands (a–c) were identified by PMF. Band (b) is the Sfibril, predicted to be a 58.7-kDa protein from amino acid sequence. (C) Phase-contrast cell images. Boxed areas are magnified at the bottom. Right: Definition of helical dimensions for helix pitch, helix width, and axial length. (D) Upper: Axial cell length of various strains (n = 25–71 cells) with and without Sfibril. Lower: Ratio of helical cells with and without Sfibril. The cells with “helix” width ≥ 0.3 µm were judged as “helical”. More than 20 field images were analyzed per strain and plotted as a data point. Statistical comparisons with and without Sfibril were evaluated by two-tailed Mann–Whitney *U* test (* *p* < 0.05, ** *p* < 0.01, *** *p* < 0.001; ns, not significant). The proteins expressed in syn3 strains are shown on the top and bottom. Data from *S. eriocheiris* are shown in rightmost graphs.

Phase-contrast microscopy showed that the cells expressing Sfibril were elongated, unlike the original syn3 cells, and 3.6% of the cells were helical (Fig. 1C, D), consistent with previous results (6, 24). In previous studies, expression of isoforms of SMreB induced syn3 cell elongation and helix formation (23, 24, 27, 30). Next, we examined the effect of additionally expressing Sfibril in cells expressing SMreB4, SMreB5, or both (SMreB45 for short), termed SMreB4F, SMreB5F, and SMreB45F, respectively (Fig. 1D). The Sfibril expression promoted cell elongation in cells with SMreB4, but not in cells with SMreB5 and SMreB45. The helix formation was promoted in cells with SMreB4 and SMreB45, but not in cells with SMreB5.

These results indicate that Sfibril can induce cell elongation and helix formation independently of SMreBs. These effects were not additive to those of single SMreBs. However, in syn3 cells with SMreB45, Sfibril markedly increased the proportion of helical cells.

### Subcellular localization and filament structure of Sfibril

To examine the localization of Sfibril, we inserted mCherry, a red fluorescent protein, after residues D81, D122, and G162 (Fig. 2A, S1, S2A, B). The cells expressing the labeled Sfibril showed elongation and helix formation with ratios of 6.7%, 11.9%, and 4.1% (elongation) and 1.8%, 3.9%, and 1.3% (helix formation) for D81, D122, and G162, respectively, comparable to those of the unlabeled Sfibril (5.1% elongation and 3.6% helicity; Fig. S2D). Sfibril fused with mCherry at D81 showed the helical dimensions closer to those of unlabeled Sfibril than other fusions (Fig. S2D). The fluorescence signal was distributed throughout the elongated cell body (Fig. 2A). The Sfibril fusion at residues D122 and G162 exhibited localization patterns similar to the fusion at D81 (Fig. 2A, Fig. S2B).

**FIG 2.**
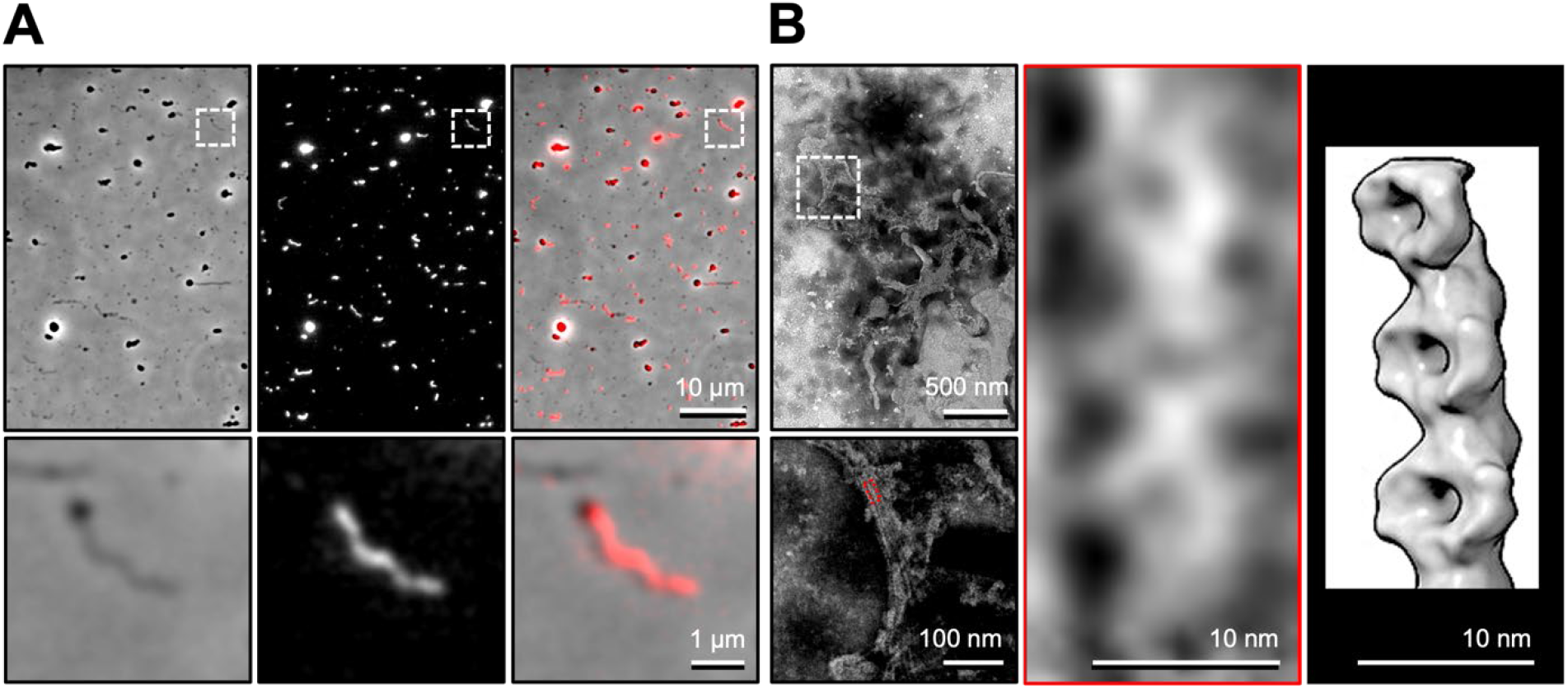
Sfibril in syn3 cells. (A) Fluorescence microscopy of syn3 cells. Sfibril was fused with mCherry at the C-terminus of residue D81 with flanking tetrapeptide linkers (SGSS at the N-terminus and SGAP at the C-terminus of mCherry) (43–46) (pSeN601; Sfibril^D81::mC^, Fig. S2) and expressed in syn3. Phase-contrast, fluorescence and merged images are shown from the left. Boxed areas are magnified at the bottom. (B) Negative-staining EM of Sfibril filaments, which were partially isolated from cell lysate. Sequential magnified views of boxed regions show connected ring-like structures similar to the three-dimensional surface structure of the isolated filament shown at the right (18).

Next, we observed the cell lysate by negative-staining electron microscopy (EM) (Fig. 2B). We found filamentous structures with chained ring units resembling those of Sfibril filaments isolated from *Spiroplasma* ribbon (Fig. 2B) (18). These results indicate that Sfibril forms filaments in syn3 and induces cell elongation and helicity formation.

### Effect of Sfibril on cell elongation

To assess the contribution of Sfibril to the robustness of cell elongation, we analyzed cell dynamics under osmotic shifts (Fig. 3 and Movies S1 and S2). Cells equilibrated in iso-osmotic PBS-G (∼310 mOsm/kg) were exposed to pure water (0 mOsm/kg) for hypo-osmotic downshift and then returned to PBS-G for recovery (Fig. 3A, B). Because cell shrinkage and recovery completed within 40 s after each buffer exchange, we compared the recovery ratio (L_rec_/L_pre_) of elongated cells (L_pre_ > 1.0 µm) across strains with and without Sfibril (Fig. 3B).

**FIG 3.**
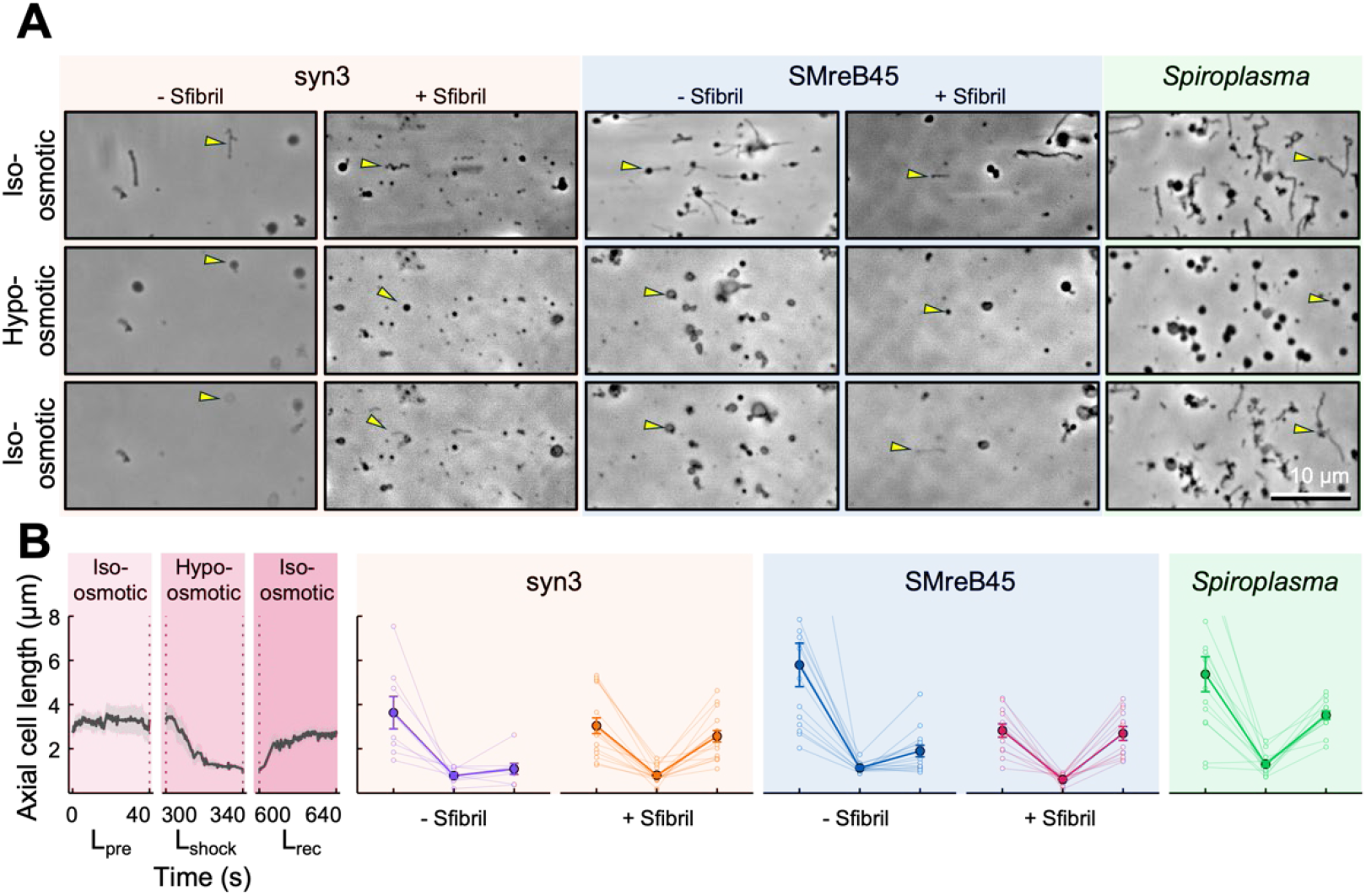
Effects of Sfibril on recovery of cell elongation from osmotic shifts. (A) Phase-contrast microscopy of cells through osmotic shifts: iso-osmotic baseline in PBS-G: 75 mM sodium phosphate, 68 mM NaCl, pH 7.3, 20 mM glucose (∼310 mOsm/kg), osmotic downshift (pure deionized water, 0 mOsm/kg), and recovery upshift (return to PBS-G). Cells were incubated under each condition for 300 s; because cell shrinkage and recovery were complete within 40 s, the initial 40-s videos were analyzed. Proteins expressed are shown on the top. *Spiroplasma* was also analyzed. Tracked cells are marked by a yellow triangle. (B) Change in axial cell length through osmotic shifts in elongated cells (L_pre_ > 1.0 µm). The leftmost panel shows the time course of cell length changes in SMreB45F (n = 12) (Movie S1, S2). The right panels show cell lengths measured at the end of each transition window: baseline (L_pre_, 40 s), maximum shrinkage (L_shock_, 340 s), and after recovery in PBS-G (L_rec_, 640 s). Faint and bold lines represent individual cells and mean ± SEM (n = 8–16 cells per strain). Within-strain L_rec_ vs. L_pre_ was evaluated using a two-tailed paired t-test (syn3, *p* = 0.018; Sfibril, *p* = 0.31; SMreB45, *p* = 8.4 × 10^-4^; SMreB45F, *p* = 0.74; *Spiroplasma*, *p* = 0.032; *p* > 0.05 indicates no statistically significant difference between recovered and baseline lengths). Between-strain recovery ratios (L_rec_/L_pre_; medians: syn3, 0.29; Sfibril, 0.86; SMreB45, 0.33; SMreB45F, 0.92; *Spiroplasma*, 0.67) were compared using the Kruskal–Wallis test (*p* = 1.5 × 10^-4^) followed by pairwise Mann–Whitney U tests (SMreB45F vs. SMreB45, Holm-adjusted *p* = 0.0013; Sfibril vs. syn3, raw *p* = 0.0071; Holm-adjusted *p* = 0.0082; *Spiroplasma* vs. SMreB45, Holm-adjusted *p* = 0.0082).

In syn3 cells, expression of Sfibril alone significantly increased the length recovery ratio compared with syn3 cells without Sfibril (Holm-adjusted *p* = 0.0082, Mann–Whitney *U* test; Fig. 3B). Similarly, co-expression of Sfibril with SMreB45 (SMreB45F) significantly increased the recovery ratio compared with SMreB45 alone (Holm-adjusted *p* = 0.0013, Mann–Whitney *U* test), restoring length recovery to a level at least as high as that of *Spiroplasma*. Recovery was also observed in syn3 cells expressing fluorescently labeled Sfibril, which localized along the entire cell axis (Fig. S3). These results indicate that Sfibril helps maintain the elongated cell shape under osmotic perturbation.

### Effects of Sfibril on swimming motility caused by SMreB45

To examine how Sfibril influences the cell motility driven by SMreB45, we tracked centroid trajectories of the syn3 cells with and without Sfibril over 9.83 s (Fig. 4A and Movie S3). The trajectories suggested that Sfibril can promote net cell translocation (Fig. 4A). The whole-body translocation was quantified using the time-averaged mean squared displacement (MSD) evaluated up to half the recording duration (Δt = 5 s). The analysis revealed a significant enhancement in translocation efficiency by Sfibril, raising MSD at Δt = 5 s from 4.29 µm^2^ to 9.38 µm^2^ (*p* = 2.66 × 10^-4^, Mann–Whitney *U* test; Fig. 4B), although this average remained far below the 600 µm^2^ observed at the same Δt for *Spiroplasma*. These findings suggest that Sfibril elevates translocation efficiency.

**FIG 4.**
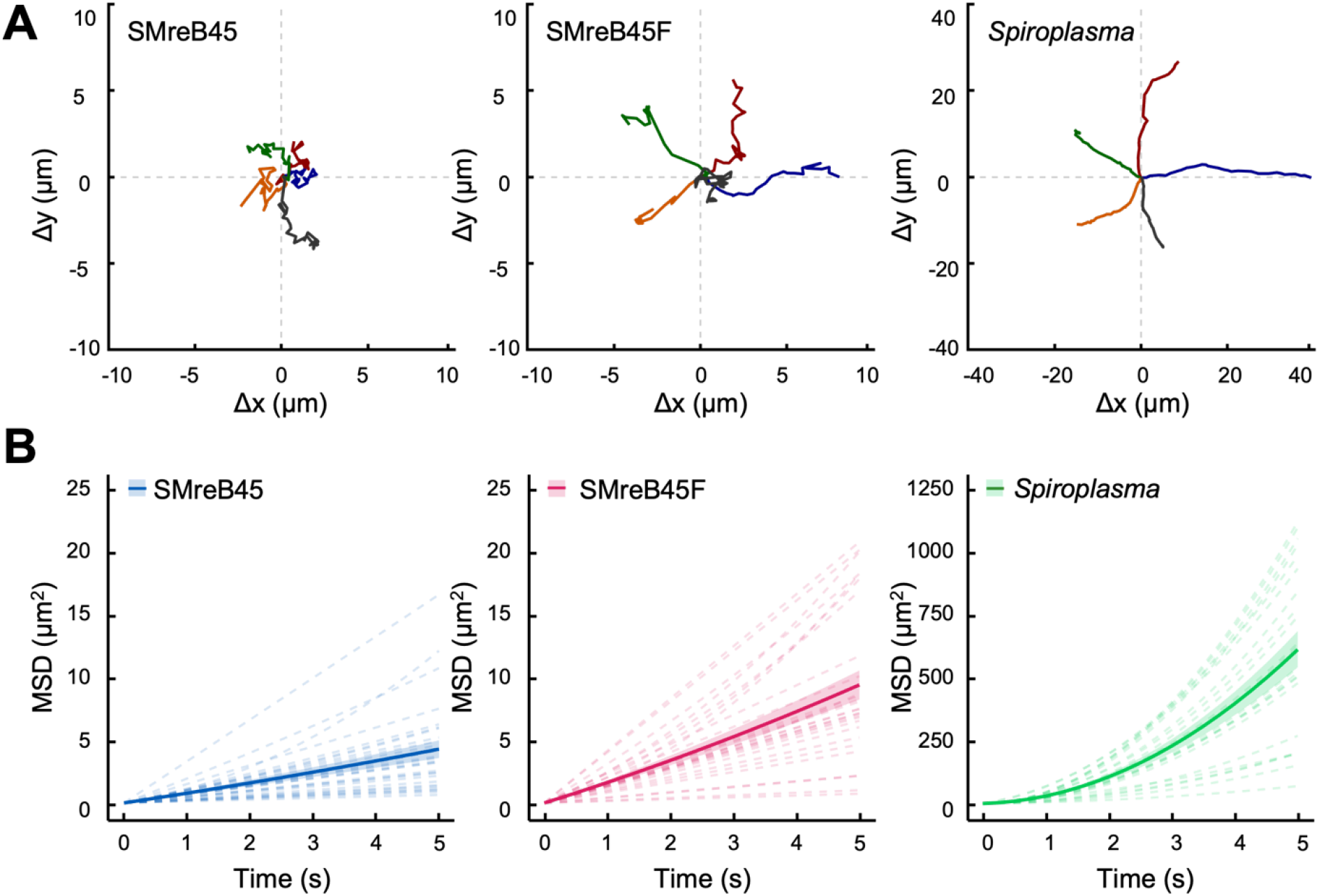
Effect of Sfibril on cell translocation. (A) Swimming trajectories of the cell centroid over 9.83 s for five cells with SMreB45, SMreB45F, and *Spiroplasma* (Movie S3). Note, different scales are applied to syn3 and *S. eriocheiris*. (B) Time-averaged MSD analysis of cell centroid up to a lag time of 4.95 s (∼5 s; 15 intervals at 0.33-s steps, representing half of the 9.83-s trajectory). Dashed lines represent individual cell fits. Solid lines with shaded areas represent the mean ± SEM (n = 22–28 cells per strain from three independent experiments). Note, different scales are applied to syn3 and *S. eriocheiris*. Statistical comparisons at Δt = 5 s were calculated using two-tailed Mann–Whitney *U* test (*p* = 2.66 × 10^-4^; \*\*\**p* < 0.001).

### Effects of Sfibril on helicity shift processivity and cell stiffness

Next, we examined the processivity of helicity shift along the cell body (Fig. 5, Fig. S4). Cell helicity handedness was determined from the optical pattern at the top plane relative to the glass surface (Fig. 5A) (31). Based on the video frames, we traced the continuous propagation of the kink (the boundary between RH- and LH-segments) along the straightened cell axis and measured the traveling distance (Fig. 5B, C, Fig. S4). Kink traveling distance was significantly increased by Sfibril expression (Fig. 5C), although the overall processivity still remained lower than that of *Spiroplasma* (Fig. 5B, C).

**FIG 5.**
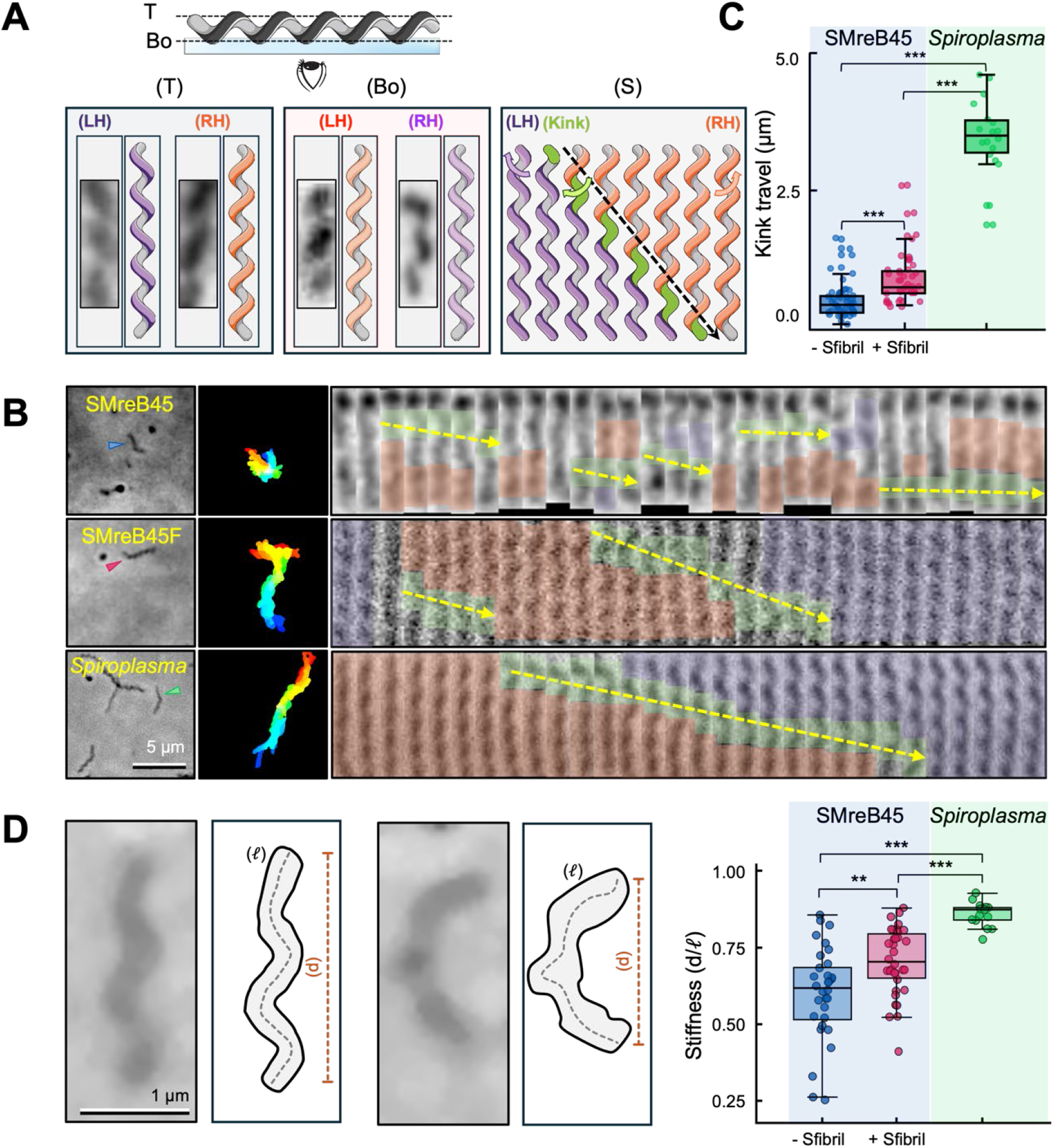
Effects of Sfibril on processivity of helicity shift and cell body stiffness. (A) Schematic illustration of cell helicity judgement. (Upper) Focal plane positioning at top (T) and bottom (Bo) relative to the glass. (T, Bo) Optical phase-contrast images and corresponding structural models for LH (purple) and RH (orange). (S) Schematic of helicity shift with kink (green), RH (orange), and LH (purple) segments traveling from upper to lower. (B) Processivity of helicity shift. (Left) Phase-contrast image at time 0 focusing on a cell. (Middle) Cumulative cell images over 10 s colored from red to blue. (Right) Montage of 30 consecutive digitally straightened cell images over 1 s (30 fps) along the 4 µm cell length. LH, RH, kink, and ambiguous regions are colored purple, orange, green, and grey, respectively. Dashed yellow arrows indicate kink movement. For kinks that persisted until the end of the video, traveling distance was measured only up to the end of the recording. (C) Kink traveling distance. Box plots show the distribution with individual data points (n = 20–40 events per strain). Statistical comparisons were evaluated using two-tailed Mann–Whitney *U* tests (***, *p* < 0.001). (D) Cell stiffness expressed as the ratio of the end-to-end distance between cell poles (d) and total cell length (ℓ). Representative elongated (left) and curved (right) syn3 cells are shown with a schematic. Averages for video frames of 1 s (30 fps) for 15–35 cells are shown (Movie S4). Significance was judged by two-tailed Mann–Whitney *U* tests (* *p* < 0.05, ** *p* < 0.01, *** *p* < 0.001).

To investigate the cause of the increased MSD, we also analyzed the cell stiffness expressed as the ratio of the end-to-end distance between cell poles (d) to the total cell length (ℓ) (9). The results indicate that Sfibril increases d/ℓ, showing a contribution to cell stiffness resisting body-bending forces (Fig. 5D, Movie S4). The increased stiffness may allow smoother helicity shifts and more efficient water propulsion.

### Effects of Sfibril on cell helicity handedness

Next, we evaluated the contribution of Sfibril on cell handedness (Fig. 6). Cells expressing Sfibril or SMreB4 with Sfibril exhibited 100% LH geometry, whereas cells expressing SMreB5 alone were 100% RH (Fig. 6A, B). Co-expression of Sfibril with SMreB5 yielded a mixed population, with 66.7% RH and 33.3% LH (Fig. 6B left). Most cells with SMreB45 showed continuous helicity shifts regardless of Sfibril (Fig. 5B). The fraction of moving cells was much larger with Sfibril than that without Sfibril (Fig. 6B right). This increase is probably related to the large proportion of helical cells caused by Sfibril (Fig. 1D), because the movement depends on the helical structure. These results may suggest that right-handedness is caused by SMreB5 and left-handedness is caused by Sfibril.

**FIG 6.**
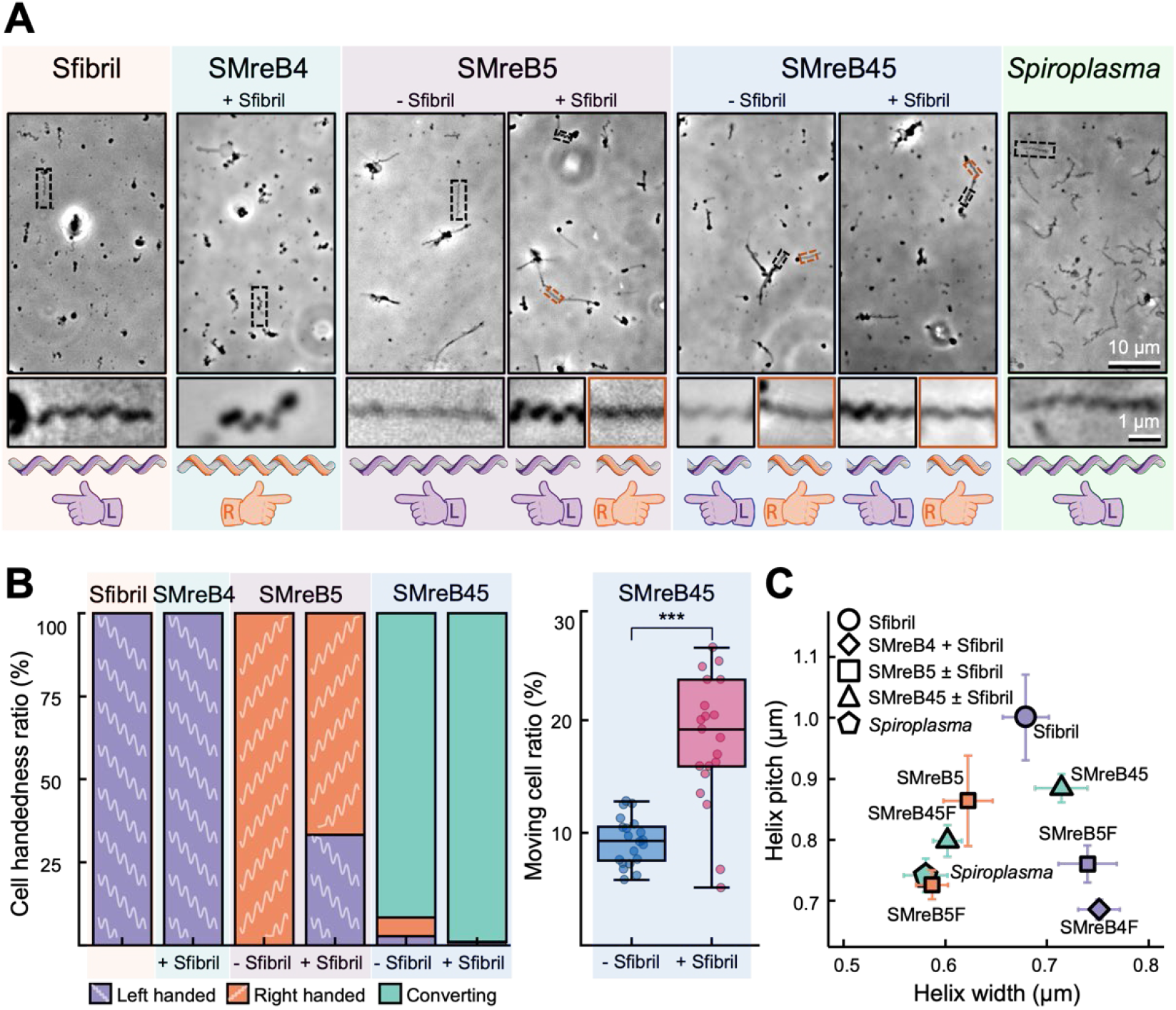
Effect of Sfibril on cell helicity handedness. (A) Phase-contrast microscopy of syn3 cells. Proteins expressed are shown on the top. *S. eriocheiris* was also analyzed. Boxed regions are magnified at the bottom. Handedness of magnified cells is illustrated at the bottom with left (L, purple) and right (R, orange) colored hand symbols, respectively. (B) Left: Cell ratios of LH (purple), RH (orange), and converting (green) (n = 20–30 cells per strain). Right: Ratio of moving cells expressing SMreB45 with and without Sfibril. The significance was judged by two-tailed Mann–Whitney *U* test (***, *p* < 0.001). (C) Helical geometry (helix pitch and helix width) of various strains. Data represent mean ± SEM (n = 30 cells per strain). Handedness is shown by colors common with those in (B).

To determine how these proteins contribute to handedness, we compared the helical geometry across different strains (Fig. 6C). We colored the data points to show their handedness. The helix width tended to be larger in LH cells than in RH cells, whereas no such tendency was observed for helix pitch. We could not measure the dimensions of LH and RH parts separately for converting cells, because their shapes changed substantially and rapidly.

## DISCUSSION

Genes encoding Sfibril are found in the genomes of most *Spiroplasma* species, and the proteins may account for nearly half of the ribbon structure that serves as motility machinery. However, the formation of the helix necessary for swimming and its helicity shifts are driven by SMreBs. Therefore, the role of Sfibril in *Spiroplasma* cells remained unknown. Here, detailed analysis based on syn3 showed that Sfibril contributes to the formation of a ribbon, the stability of cell morphology, and the efficiency of swimming. These results are consistent with previous studies (5, 6).

### Role for cell elongation and helix formation

The effect on the structural stability required for swimming was particularly evident in the syn3 strain, which possesses motility by SMreB45. With the expression of Sfibril, the proportion of cells forming helix increased from 2.3% to 17.4% (Fig. 1D), and the ratio of moving helical cells increased from 9.3% to 19.2% of the total cell population (Fig. 6B). Furthermore, the role of Sfibril in structural stability was also evident in syn3’s recovery from osmotic shock (Fig. 3, S3). These roles are related to the fact that isolated Sfibril filaments are highly stable and cannot be broken down into anything smaller than single-stranded filaments, even by intense ultrasonic treatment (18, 19). This robustness contrasts with MreB filaments, which depolymerize easily (27, 30).

### Role for processive swimming

Analysis of swimming behavior showed that Sfibril significantly increased the distance traveled from the starting point (Fig. 4). One contributing mechanism may be the more continuous occurrence of helicity shifts, the force-generation process responsible for swimming (Fig. 5, S4). This may be because Sfibril suppresses cell bending at large angles: when SMreB filaments—which are inherently nearly linear—are significantly bent by an external force, the free conformational changes in the bent region may be impeded (Fig. 5D).

### Role for helicity shifts

Another role of Sfibril in swimming may be to assist the helicity shift itself (Fig. 6). SMreB5 forms an RH helix, and when SMreB4 is present, it occasionally shifts to an LH helix. This may suggest that the LH Sfibril guides the shift of the RH SMreB5 into an LH helix. In other words, Sfibril may allow the SMreB4 and SMreB5 pair to shift their helicity with a lower energy barrier. This idea is further supported by the observation that the presence of Sfibril increased the ratio of moving cells (Fig. 6B).

When the filaments composed of SMreB4 and SMreB5 actively shift helicity, does the handedness of Sfibril also shift? All syn3 helical cells that expressed only Sfibril showed left-handedness. In addition, all of the isolated Sfibril filaments are LH (18). These facts suggest the following two possibilities. One is that the helicity shift of Sfibril is caused by direct force from SMreBs. It has been reported that Sfibril binds to the SMreBs (23, 32). Moreover, the helical geometry of syn3 with Sfibril was different from syn3 cells additionally expressing SMreB5 or SMreB4, even though they formed LH helices (Fig. 6C). Another possibility is that Sfibril filaments remain LH, and the cell’s helicity is determined by the SMreB filaments.

In this study, the helical structure and swimming ability of syn3, reconstituted with SMreB4 and SMreB5, showed marked improvements in robustness and efficiency upon the addition of Sfibril. However, *Spiroplasma* cells demonstrated far higher swimming efficiency and kink processivity. This difference may arise from other ribbon components, SMreB1, SMreB2, and SMreB3, and the complete organization of the ribbon.

## MATERIALS AND METHODS

### Bacterial cultures

JCVI-syn3B (GenBank CP069345.1) and *S. eriocheiris* (TDA-040725-5) were cultured statically in liquid SP4 medium containing 50 µg/mL ampicillin at 37°C and 30°C, respectively, to an OD_620_ of 0.03–0.05 (24). syn3 transformants were selected with 3 µg/mL puromycin. *Escherichia coli* DH5α was used for plasmid preparation.

### Strain construction

Plasmids were constructed from pSeW001 (24) (Fig. S1). Amplified DNA fragments were assembled with the plasmid fragment using NEBuilder HiFi DNA Assembly Master Mix (New England Biolabs, Ipswich, MA, USA), cloned with *E. coli* DH5α (33) and introduced into syn3 cells (34). Genomic integration was verified by PCR and Sanger sequencing. Structural models of Sfibril and mCherry-fused Sfibril filaments were predicted by AlphaFold3 (35) and analyzed by UCSF ChimeraX (36). Protein expression profiles were analyzed by SDS-PAGE and mass spectrometry as previously described (18, 24).

### Microscopy

Cultured cells were harvested by centrifugation at 9,000 × g for 10 min, resuspended in iso-osmotic PBS-G (75 mM sodium phosphate, 68 mM NaCl, 20 mM D-glucose, pH 7.3; ∼310 mOsm/kg, calculated from the solute concentrations) to a 10-fold higher cell density. The cell suspension was loaded into flow chambers constructed with double-sided tape and coverslips.

For motility assay, the PBS-G in a chamber was replaced with PBS-G containing 0.5% (w/v) methyl cellulose. Both syn3 and *S. eriocheiris* were examined under the same buffer and chamber conditions. Cells were observed under an inverted microscope IX71 (Olympus, Tokyo, Japan) equipped with a UPlanSApo 100×/1.40 NA Ph3 objective. For fluorescence imaging, the microscope was equipped with a Texas Red filter set (TXRED-4040B-OMF; Semrock, Rochester, NY, USA). Images and videos were recorded with a complementary metal-oxide semiconductor (CMOS) camera, DMK33UX174 (The Imaging Source Asia Co., Ltd., Taipei, Taiwan), operating at 30 frames per second (fps).

For the osmotic shift assay, the collected cells were suspended and kept in PBS-G for 300 s in a flow chamber to adhere spontaneously to the glass; as the cells attached to the glass were analyzed, the effects of osmolality on binding cannot be excluded. Hypo-osmotic downshift was performed at 300 s by replacing the chamber solution with pure water (0 mOsm/kg) and kept for 300 s, after which the buffer was returned to PBS-G at 600 s and kept for 300 s for recovery. Cell lengths were analyzed during the initial 40 s of each 300-s window (0-40 s for baseline, 300–340 s for hypo-osmotic shock, and 600–640 s for recovery), within which osmotic shrinkage and recovery reached steady states. Negative-staining EM and cell extraction were conducted as previously described (18).

### Video analyses

All image processing and video analyses were performed using Fiji (37) and its plugins as follows. Cell centroids in phase-contrast videos recorded at 30 fps were tracked using MTrackJ (38). MSD was calculated at 0.33-s lag-time intervals up to 4.95 s (∼5 s, intentionally restricted to the first half of the 9.83-s trajectories to ensure reliable statistical sampling across lag intervals) (39). Actively moving helical cells were distinguished from stationary or freely diffusing cells by time-lapse trajectory visualization using Color Footprint (40) as described previously (29). Cell helicity handedness was judged at upper and lower focal planes relative to the coverslip (31). Helicity shift was tracked by straightening cell axes (24). Kink traveling distance was measured as the axial displacement from the initiation to the termination of kink propagation. Helical dimensions (helix pitch and helix width) were measured as previously described (Fig. 1C) (24). For osmotic shift analysis, elongated cells with an initial baseline length L_pre_ > 1.0 µm (exceeding the ∼0.5–0.8 µm diameter of spherical parent syn3 cells; Fig. 1D) were selected so that axial shrinkage and re-elongation could be reliably resolved, and their axial lengths were measured using JFilament (41). To determine cell centroids (Movie S3) and quantify body stiffness (Movie S4), phase-contrast images were binarized by intensity thresholding, thinned to one-pixel-wide cell centerlines (binarized cell axes), and measured for contour length (ℓ) and end-to-end distance between terminal endpoints (cell poles; d) using the Skeletonize and Analyze Skeleton (2D/3D) plugins (42). Cell stiffness was evaluated as the ratio d/ℓ averaged over 30 consecutive frames (1 s) (9).

### Statistical analysis

Statistical analyses and graphical plots were performed using R (R Foundation for Statistical Computing) and Python. Differences between two groups (including planned pairwise comparisons with and without Sfibril in Fig. 1D, 5C, 5D, and 6B) were evaluated by two-tailed Mann–Whitney *U* tests or two-tailed paired Student’s *t*-tests, and multi-group comparisons in Fig. 3B were evaluated by Kruskal–Wallis tests followed by Holm-adjusted Mann–Whitney *U* tests across three planned pairwise comparisons (SMreB45F vs. SMreB45, Sfibril vs. syn3, and Spiroplasma vs. SMreB45) or one-way ANOVA followed by Tukey’s HSD post hoc test (Fig. S2D), as indicated in the figure legends. Data were pooled from three independent cultures (biological replicates) harvested at the same growth phase (OD620 = 0.03–0.05) that exhibited consistent trends for each strain. *p* < 0.05 was considered statistically significant. *n* indicates the number of cells analyzed (or events/fields, as specified in the legends).

## Supporting information

Supplemental file and will be used for the link to the file on the preprint.

## ACKNOWLEDGMENTS

We thank T. Shimonaka at Osaka Metropolitan University for the technical assistance with mass spectrometry. This study was supported by JST CREST and ARiSE grants (JPMJCR19S5, JPMJAR2609), Grant-in-Aid for Scientific Research (B) (JP26K01981), and a grant from the Institute for Fermentation, Osaka (IFO) (G-2026-2-111) to MM. HK is a recipient of a JSPS Research Fellowship (JP24KJ0189).

## AUTHOR CONTRIBUTIONS

A.A., H.K., and M.M. designed the research; A.A., H.K., Y.O.T., and Y.S. performed the experiments and analyzed the data; A.A. and M.M. wrote the manuscript.

## DATA AVAILABILITY

All data supporting the findings of this study are available within the paper and its supplemental material, and raw microscopy videos and tracking datasets are available from the corresponding author upon reasonable request.

