## Supplemental file and will be used for the link to the file on the preprint. for "Reconstitution in JCVI-syn3B suggests the roles of the *Spiroplasma* Fibril protein (Sfibril) in swimming": 20261117_Fibril_bioRxiv_supp_AA.pdf

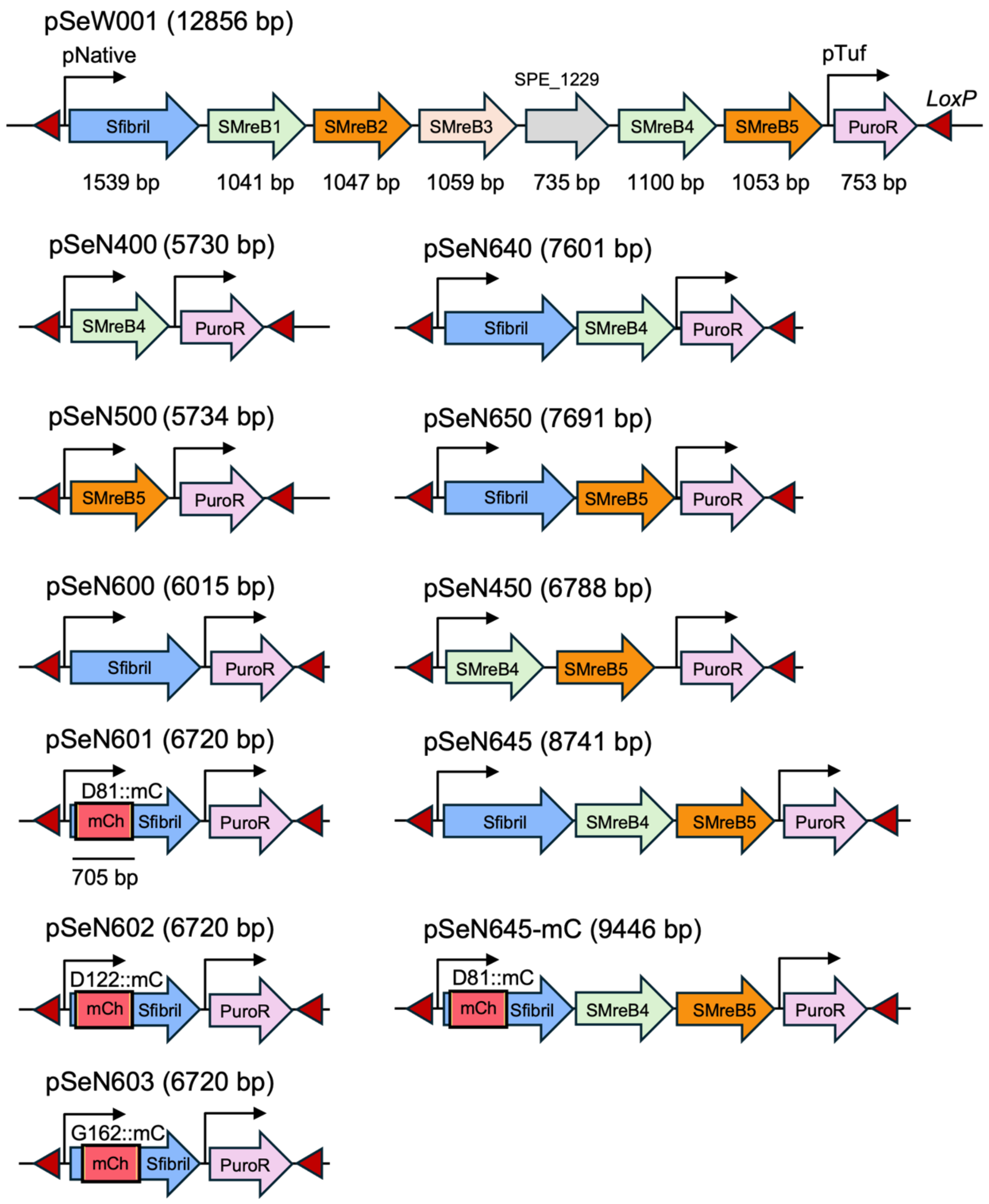


**FIG S1** DNA fragments transferred into *loxP* sites of syn3 genome. The DNA constructs used in this study were prepared from a plasmid pSeW001 coding eight genes between two *loxP* sites (24). Promoter sequences are shown as an arrow. The plasmid name and size are shown above the schematics.

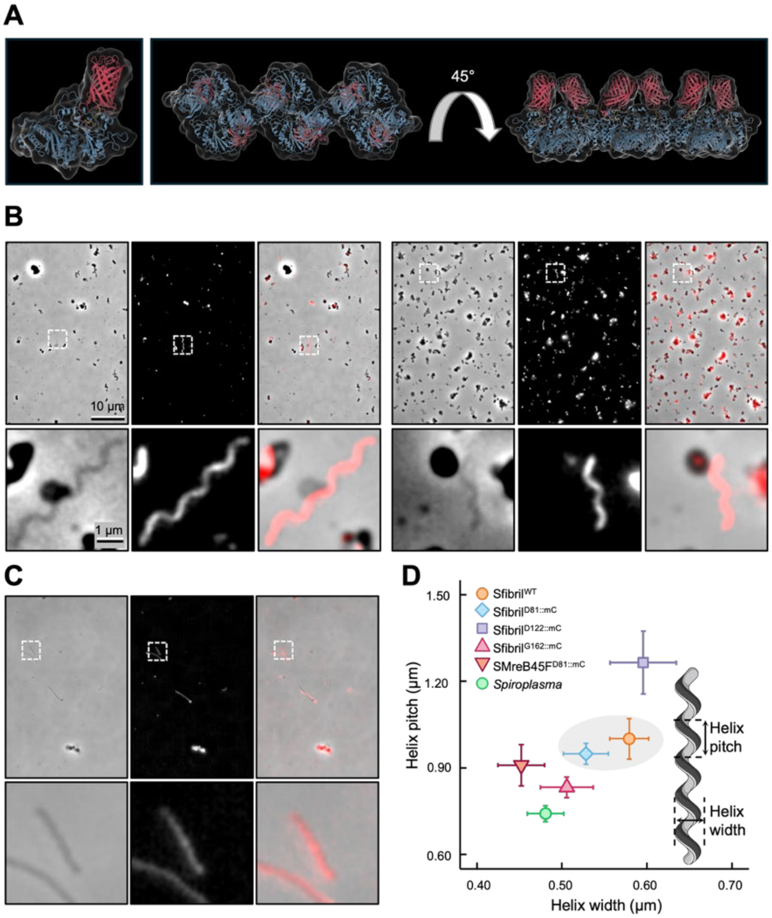


**FIG S2** syn3 cells expressing mCherry-fused Sfibril. (A) Structural model of Sfibril (blue) fused with mCherry (red) at the C-terminus of D81 flanked by flexible linkers (yellow), named as Sfibril$^{D81::mC}$. Monomer (left) and filament (right) are shown. The fluorescence cell images are shown as Fig. 2. (B) Fluorescence cell images expressing Sfibril fused with mCherry at G123 (left) and G162 (right). (C) Fluorescence images of cells expressing SMreB45 and Sfibril$^{D81::mC}$. (B, C) Phase-contrast, fluorescence, and merged images are shown from the left. Boxed areas are magnified at the bottom. (D) Helical geometry of various cells shown as mean ± SEM (n = 30 cells per strain). Data for cells expressing original Sfibril and Sfibril$^{D81::mC}$ are circled gray. Helical dimensions were evaluated using one-way ANOVA followed by Tukey's HSD post-hoc test.

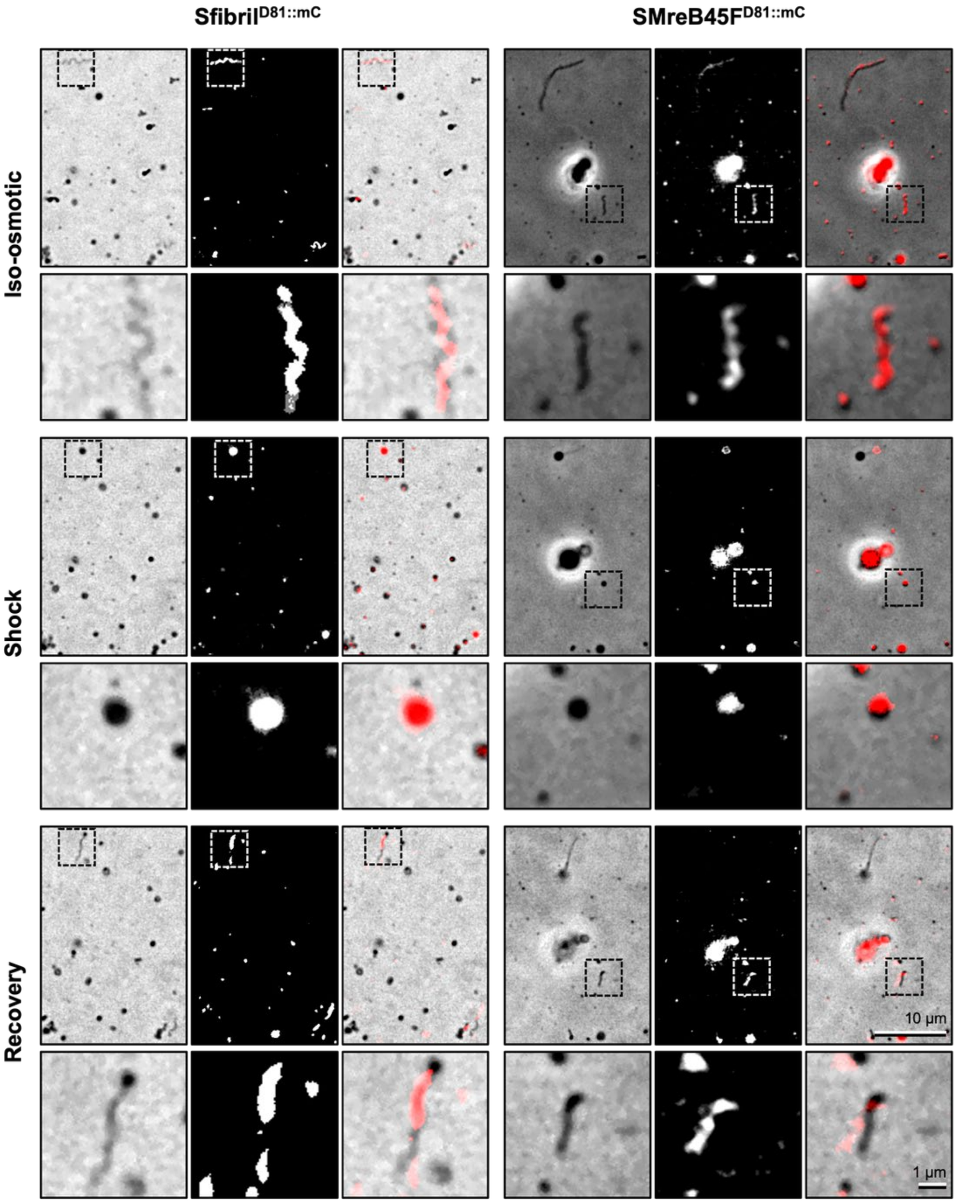


**FIG S3** Cell shape transformation under osmotic shifts in cells with fluorescent Sfibrils. Phase-contrast, fluorescence, and merged images during osmotic shifts are shown from the left. Boxed areas are magnified at the bottom. Fluorescence cell images expressing Sfibril fused with mCherry at residue D81 alone (Sfibril$^{D81::mC}$, left) and together with SMreB45 (SMreB45 + Sfibril$^{D81::mC,}$ right). See Fig. 2 and Fig. 3 for details.

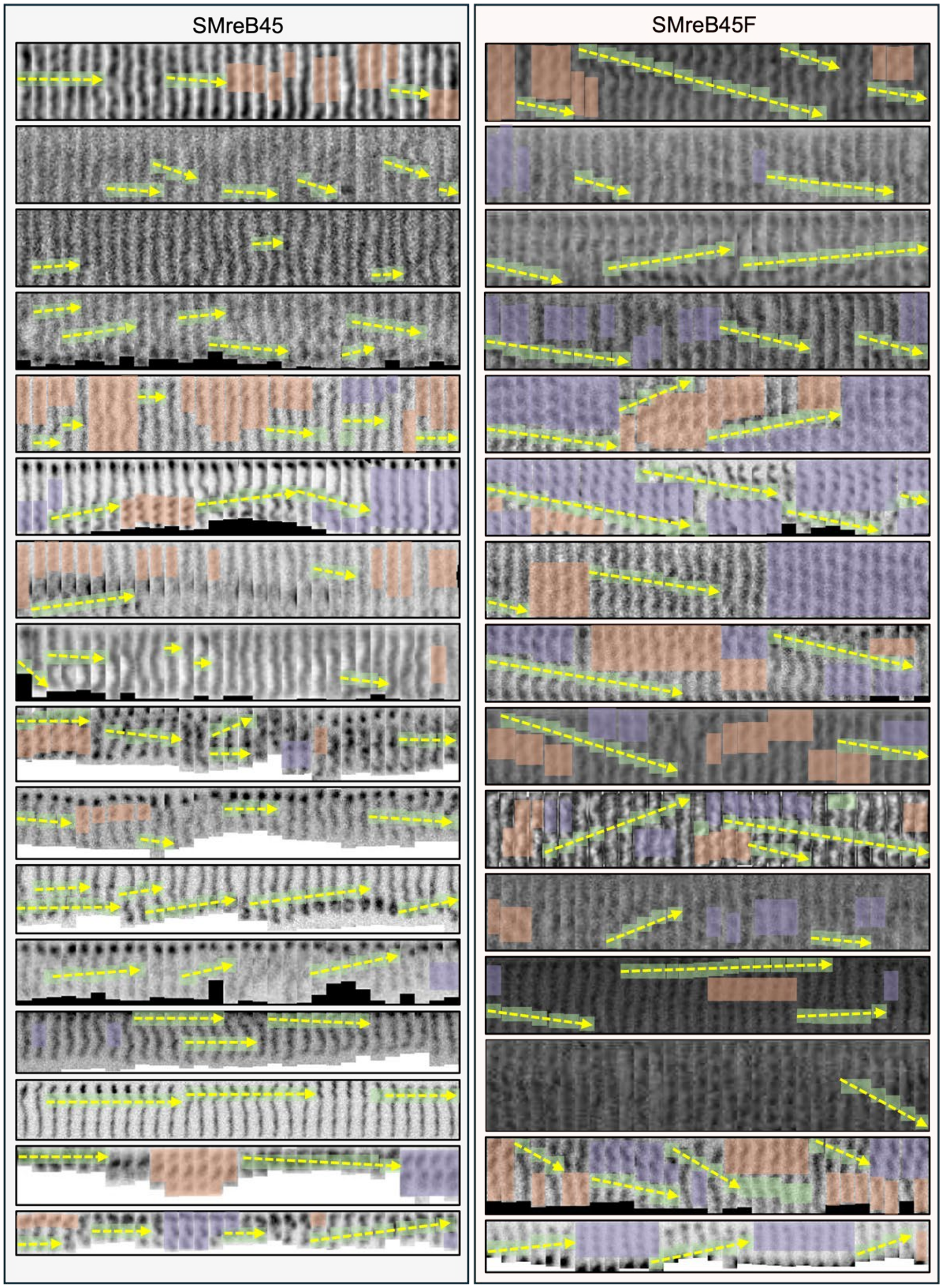


**FIG S4** Montage of helicity shifting syn3 cells. Consecutive 30 straightened cell images are shown over 1 s (30 fps) for 15 cells of SMreB45 (left) and SMreB45F (right). See Fig. 5 for more details.

**Movie S1** Cell shape changes through osmotic shifts. Cells were incubated for 300 s in each condition (iso-osmotic baseline, hypo-osmotic downshift, and recovery upshift); the initial 40 s of each transition is shown.

**Movie S2** Cell shape changes in 180 s after osmotic shifts. Eight representative cells are shown for syn3 expressing SMreB45F (left) and *Spiroplasma* (right).

**Movie S3** Centroid trajectories of moving cells. (Left) Phase-contrast images (upper) and one-pixel-wide cell centerline (lower). Cell axis, centroid, and centroid trajectories over 9.83 s are colored white, red, and rainbow from yellow to blue, respectively. (Right) Integrated centroid trajectories of five cells each with SMreB45 (blue) and SMreB45F (red), aligned to a common start point.

**Movie S4** Cell stiffness analysis. Phase-contrast (left) and binarized cell axes with cell poles (yellow and magenta circles) (right) are shown for SMreB45 (left), SMreB45F (middle), and *Spiroplasma* (right) cells over 1.00 s (30 fps).

**Table S1** Identification of proteins from SDS-PAGE bands by peptide mass fingerprinting (PMF)

| **Band** | **Gene ID** | **Protein Name** | **Mascot Score[1]** | **Mass (kDa)[2]** |
|---|---|---|---|---|
| **a** | SPE_0123 | Lipoprotein | 105 | 64.4 |
| **b** | SPE_0666 | Sfibril | 249 | 58.7 |
| **c** | glpK | Glycerol kinase | 103 | 57.3 |

[1] Mascot search score; scores > 67 indicate significant identity or extensive homology ($p < 0.05$).

[2] Theoretical molecular weight calculated from the amino acid sequence.

**Table S2: List of primers**

**Table S3: List of plasmids and their sequences.**
